# The RNA binding proteins ZFP36L1 and ZFP36L2 modulate transcriptional and post-transcriptional genome-wide effects of glucocorticoids

**DOI:** 10.1101/2022.09.27.509553

**Authors:** Jennifer Rynne, Elena Ortiz-Zapater, Paniz Khooshemehri, Nicole O. Ponde, Giselda Bucca, Andrew Hesketh, Manuela Platé, Rachel Chambers, Colin Smith, Rocio T. Martinez-Nunez

## Abstract

Glucocorticoids (GCs) are one of the most used anti-inflammatory drugs worldwide. Despite their widespread use, our understanding of their post-transcriptional effects remains poorly understood. The tristetraprolin (TTP) RNA binding protein (RBP) family (ZFP36, ZFP36L1 and ZFP36L2) has been implicated in inflammation regulation via binding to AU-rich elements (ARE) in mRNAs, with TTP being implicated in GC modulation. We hypothesised that ZFP36L1 and ZFP36L2 are part of the GC pathway and tested this hypothesis in bronchial epithelium, which commonly encounters GC *in vivo* upon inhalation. Our data show that dexamethasone, a commonly used GC, modulated the levels, subcellular localisation and RNA binding of ZFP36L1/L2. Employing Frac-seq (subcellular fractionation and RNA-sequencing), we show that GC modulated distinct subsets of RNAs in a subcellular-dependent manner. In addition to their mostly known transcriptional effects (116 differentially expressed genes, DEGs), GCs modified the binding to monosomes of myriad mRNAs (83 differentially bound genes, DBGs). We also demonstrate that ZFP36L1/L2 modulated gene expression mainly at the total cytoplasmic and polyribosome binding levels. ZFP36L1/L2 down-regulation led to an increase in ARE-containing mRNAs and a pronounced modification of the effects of GC on gene expression. We observed a small overlap of genes modulated by GCs when comparing control and ZFP36L1/L2 knockdown cells, in a subcellular-dependent manner Our data also suggest a novel role for these RBPs and GCs in epithelial biology via regulation of mRNAs encoding proteins important for epithelial cell function including cellular structure. We believe that our data has further implications in how we investigate gene expression. We show the power of employing sub-cellular fractionation when analysing genome-wide effects for known ‘transcriptional modulators’ such as GCs, as well as a tool to demonstrate the extent of the effect of RBPs on gene expression modulation beyond total RNA levels.

## Introduction

Glucocorticoids (GCs) are steroid hormones involved in regulating multiple physiological processes, including metabolism and inflammation. GCs such as dexamethasone, fluticasone, budesonide, hydrocortisone or prednisolone are some of the most prescribed drugs in the world due to their potent anti-inflammatory action, with estimations from 14% (1) to 3% (2) of the adult population taking oral GCs. Respiratory diseases such as asthma account for 40% oral GC prescriptions (3). In addition to their anti-inflammatory effects, long-term use of GCs leads to multiple side effects including diabetes, obesity or depression (4). The action of GCs relies on binding to the glucocorticoid receptor (GR), which is a steroid-hormone nuclear receptor present in most human tissues (5). GR has two GR isoforms, GRα and GRβ (6) and can be phosphorylated by MAP kinases and cyclin-dependent kinases at its serine residues in a ligand-dependent and independent manner (7).

GCs are mainly known as transcriptional regulators. Binding of GCs to cytoplasmic GR induces a phosphorylation and conformational change leading to GR dissociation from its chaperones, heat shock protein (Hsp) 70, 90 and p23 (8). Subsequently, GR translocates to the nucleus where it homodimerizes and binds to DNA glucocorticoid response elements (GREs), acting as a transcription factor (9). Its activity as transcription factor depends on the GRE sequence, activity of co-factors and chromatin accessibility (10, 11). GCs can also act via protein-protein interactions, for example by repressing transcription factors such as the nuclear factor kappa B (NF-kB), activator protein 1 (AP-1) or signal transducer and activator of transcription (STAT) family (9). GCs also possess post-transcriptional effects, mostly circumscribed to AU-rich (ARE)-mediated decay via induction of mitogen-activated protein (MAP) kinase phosphatase 1 (MKP-1) and tristetraprolin (TTP) expression (12). Many genes are regulated post-transcriptionally by GCs through mRNA stability including *COX2, CCND3, CCL11, GMCSF, IFNB, IL1, IL6* or *CXCL8* (13). In addition, GR has been shown to function as an RNA binding protein itself (14, 15) and it was recently associated with glucocorticoid mediated decay (GMD), which requires a ligand, the loading of GR onto a target mRNA, upstream frameshift 1 (UPF1) and proline-rich nuclear receptor coregulatory protein 2 (PNRC2) (16). Unlike other decay pathways such as nonsense mediated decay, GMD does not appear to rely on translation (17). Additionally, GCs can act via non-genomic signalling pathways via membrane-associated receptors and second messengers. The characteristics of non-genomic effects include a short time frame, a different pharmacological profile because the effects are insensitive to transcriptional and translational inhibitors, efficacy on non-nucleated cells such as platelets, and the ability of steroid analogues that cannot enter intracellular compartments to elicit a response (13). However, despite their wide use and effects, coupling transcriptional with post-transcriptional effects of GCs remains poorly described and understood.

The TTP family is comprised of three CCCH zinc finger proteins (ZFP), namely ZFP36 (alias TTP or TIS11), ZFP36L1 (alias TIS11B and BRF1) and ZFP36L2 (alias TIS11D or BRF2) (18). Although structurally similar, these family members have distinct functions and targets. (19). The most studied member of the family, TTP, binds to AUUUA sequence repeats in the 3’ UTR (Untranslated Region) of its targets, promoting deadenylation of the poly(A) tail, RNA instability and degradation (20). ZFP36, ZFP36L1 and ZFP36L2 have different patterns of expression and effects. ZFP36L1 and ZFP36L2 can modulate mRNA levels in different manners, from ribosome association (21) to decay via deadenylation (22). These RBPs have been implicated in the regulation of B and T cell biology and immunity (21, 23–25). Changes in single nucleotide polymorphisms (SNPs) and/or in the expression of ZFP36L1 and ZFP36L2 have been associated with multiple inflammatory disorders (26). There is some evidence of GCs modulating ZFP36L1 and/or ZFP36L2 expression levels (27), but the role of these RBPs in the effects of GCs genomewide has not been characterised.

Based on the above, we hypothesised that GCs act via ZFP36L1 and ZFP36L2 and we set out to investigate their transcriptional vs post-transcriptional effects in bronchial epithelial cells. We chose this cell type because they sit at the interface between the airways and the environment in the lungs, being the main cells in central airways that GCs will encounter when inhaled. Our data show that GCs modulate the levels, subcellular localisation and transcript binding affinity of ZFP36L1 and ZFP36L2. We performed Frac-seq (subcellular fractionation and RNA-sequencing) in primary bronchial epithelial cells depleted of ZFP36L1 and ZFP36L2 and stimulated with GCs, which allowed us to examine mRNA expression levels in three subcellular fractions: total cytoplasmic (Total), monosome-bound mRNAs (80S) and polyribosome-bound mRNAs (Poly). These experiments revealed novel findings. Firstly, we found an unprecedented level of regulation of mRNA expression by GCs, not only transcriptionally but also at the level of monosome binding. In addition, we demonstrate that ZFP36L1 and ZFP36L2 modulated distinct mRNA targets in a sub-cellular dependent manner, with cytoplasmic RNA levels being the main target and secondly the polyribosome association of a distinct set of mRNAs. ZFP36L1/L2 downregulation caused an increase in ARE-containing mRNAs in both total and polyribosome compartments, as well as an over-representation of previously reported ZFP36L2-bound transcripts. Moreover, ZFP36L1/L2 depletion transformed the transcriptional and post-transcriptional effects of GC, in many cases causing opposite effects on gene expression modulation, particularly at the Total and 80S binding levels. Our data highlight the value of performing sub-cellular analysis when investigating RBP effects and reveal how GCs act via and modulate the activity of ZFP36L1 and ZFP36L2.

## Results

### ZFP36L1 and ZFP36L2 expression and localisation are modulated by glucocorticoids

We firstly set out to determine if GCs modulated the expression of ZFP36L1 and ZFP36L2 in bronchial epithelial cells. BEAS-2B cells were plated, starved overnight, stimulated with 10^−7^M dexamethasone (GC herein) and harvested at different time points. Both *ZFP36L1* and *ZFP36L2* mRNAs were significantly induced early after 2h stimulation; this was sustained at 24h for *ZFP36L2* (Figures 1A and 1B). 6h stimulation with dexamethasone significantly induced *ZFP36L2* mRNA (Figure 1B) and *ZFP36L1* mRNA showed a trend towards increased levels (Figure 1A). Glucocorticoid-induced leucine zipper (*GILZ*) mRNA expression was assessed as a positive control for GC effects, since it is a known GC target in airway epithelium (28). As expected, *GILZ* mRNA was significantly increased at all time points (Figure 1C). At the protein level, GC upregulated ZFP36L1 and ZFP36L2 at 2h (Figures 1D and 1E, respectively) with a trend towards upregulation for ZFP36L2 at 24h (Figure 1E), while GR expression significantly decreased at 24h (Figure 1F) as previously shown in HeLa cells (29). To further confirm our data, we determined the expression of *ZFP36L1* and *ZFP36L2* mRNAs in HBEC3-KT (3KT) cells, another bronchial epithelial cell line with the ability to differentiate into ciliated epithelium in air-liquid interface cultures (30). Figure 1G shows that the mRNA expression of *ZFP36L1* and *ZFP36L2* was also increased by GCs in 3KT cells at different time points.

**Figure 1.**
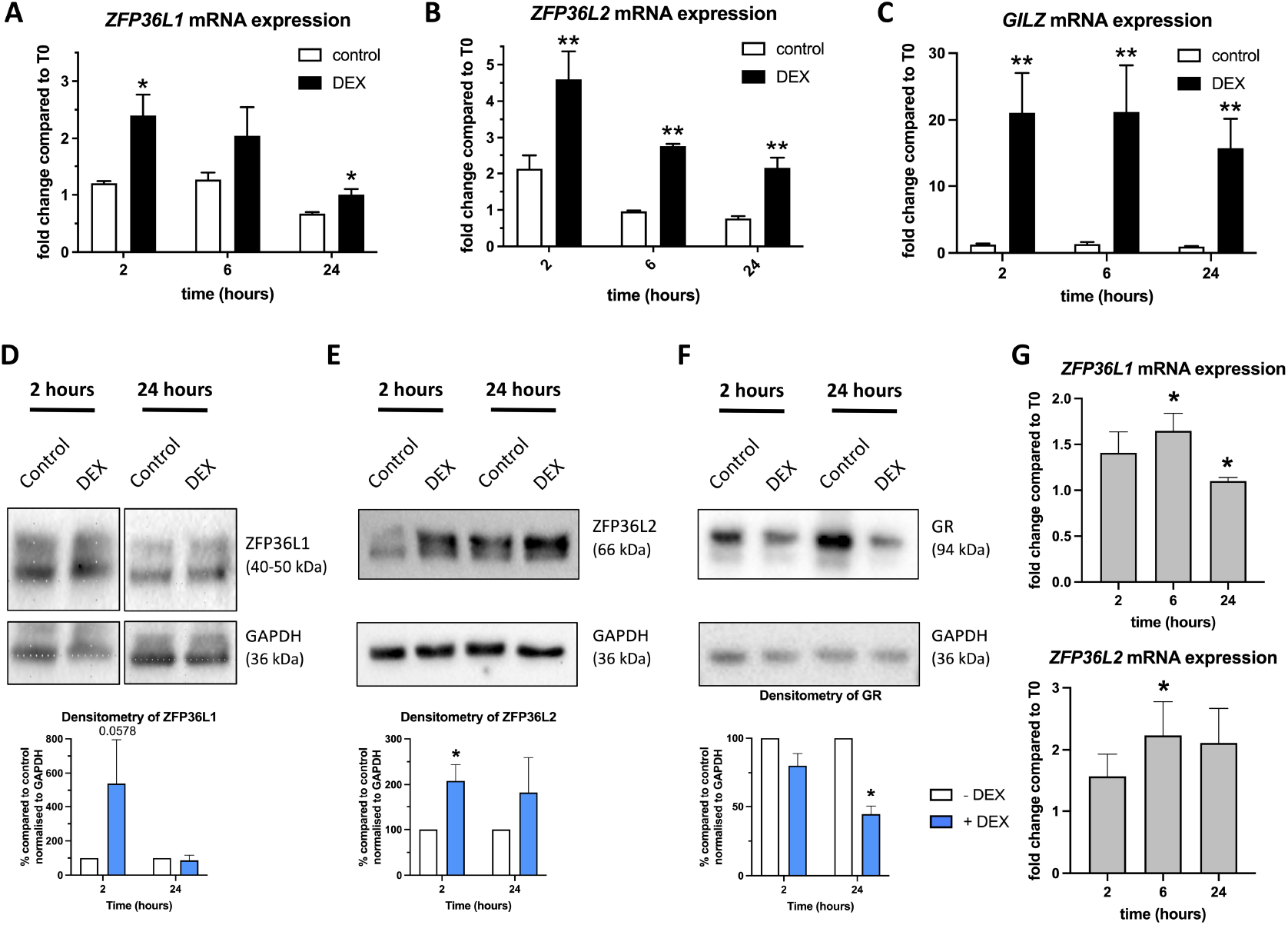
Dexamethasone modulates ZFP36L1 and ZFP36L2 expression in bronchial epithelial cells. BEAS-2B cells were stimulated with 10^−7^ M dexamethasone (DEX), harvested and analysed at different time points. Results demonstrate treatment compared to vehicle. **A-C**. Bar graphs representing mean fold change of mRNA expression (± SEM) as determined by RT-qPCR amplifying *ZFP36L1* **(A)**, *ZFP36L2* (**B**) and *GILZ* (**C**). Values were normalised to 18S. n=4. Statistical significance was assessed by two-tailed t-tests on log transformed data. **D-F**: Representative images of western blot protein analysis probing for ZFP36L1 (**D**), ZFP36L2 (**E**) and GR (**F**). Bar graphs underneath represent mean densitometry quantification of band intensity (± SEM) normalised to GAPDH. n=3-5. Statistical significance was assessed by one-tailed t-tests. **G**. Bar graphs representing mean fold change of mRNA expression (±SEM)of 3KT cells stimulated with 10^-7^ M dexamethasone of *ZFP36L1* and *ZFP36L2* normalised to 18S, n = 3-5 independent experiments. p < 0.5 *, p < 0.01 ** p < 0.001 ***.

Taken together, our data show that dexamethasone increased the expression of ZFP36L1 and ZFP36L2, suggesting a potential role for ZFP36L1 and ZFP36L2 in GC effects.

### Glucocorticoids modulate binding of ZFP36L1/L2 and GR to target mRNAs

We next investigated the potential for GCs to modulate the binding of ZFP36L1 and ZFP36L2 (ZFP36L1/L2 herein) to their targets. To this end, we employed RNA immunoprecipitation (RIP) of ZFP36L1/L2 as well as that of GR and assessed several candidate mRNAs. Rabbit IgG was used as a negative control to account for unspecific binding. We employed a dual ZFP36L1/L2 antibody (BRF1/2, Cell Signaling Technology) which has been previously shown to be selective for *Zfp36l1* in mice (25), and others have used it to immunoprecipitate ZFP36L1 in 293 T cells (22). In our hands, the dual ZFP36L1/L2 immunoprecipitated ZFP36L2 and likely ZFP36L1 in BEAS-2B cells (Supplemental Figure 1) and thus we refer to the immunoprecipitated RNA as ‘ZFP36L1/L2’ bound without distinction between the two RBPs.

BEAS-2B cells were stimulated with GC and 24h later RIP was performed. We considered ‘bound’ those mRNAs that showed statistically significant enrichment when compared to their corresponding IgG control to account for potential unspecific binding to immunoglobulin or beads. Thus, we compared ZFP36L1/L2 no GC vs IgG no GC and ZFP36L1/L2 + GC vs IgG + GC. We also took into consideration the effects of GC on the levels of the assessed target mRNAs (INPUT in Figure 2). Figure 2A shows the levels of mRNAs that were bound by ZFP36L1/L2 following a similar pattern to that of the corresponding input levels. *IL6* mRNA was decreased by GC (top panel, INPUT), while its binding to ZFP36L1/L2 showed a similar trend. *CXCL8* (*IL8* herein) transcripts did not show regulation by dexamethasone treatment (INPUT) and were similarly bound by ZFP36L1/L2 in the presence and absence of dexamethasone (Figure 2, middle graphs). *VEGF*, which has been shown to be bound by ZFP36L1 (22) also showed binding to ZFP36L1/L2 in bronchial epithelial cells, and this binding was similar with/without dexamethasone. In our hands, none of these transcripts were directly bound by GR, either in the absence or presence of GC (blue bars in Figure 2A), except for *ZFP36L1* and *ZFP36L2* mRNAs (Supplemental Figure 2).

**Figure 2.**
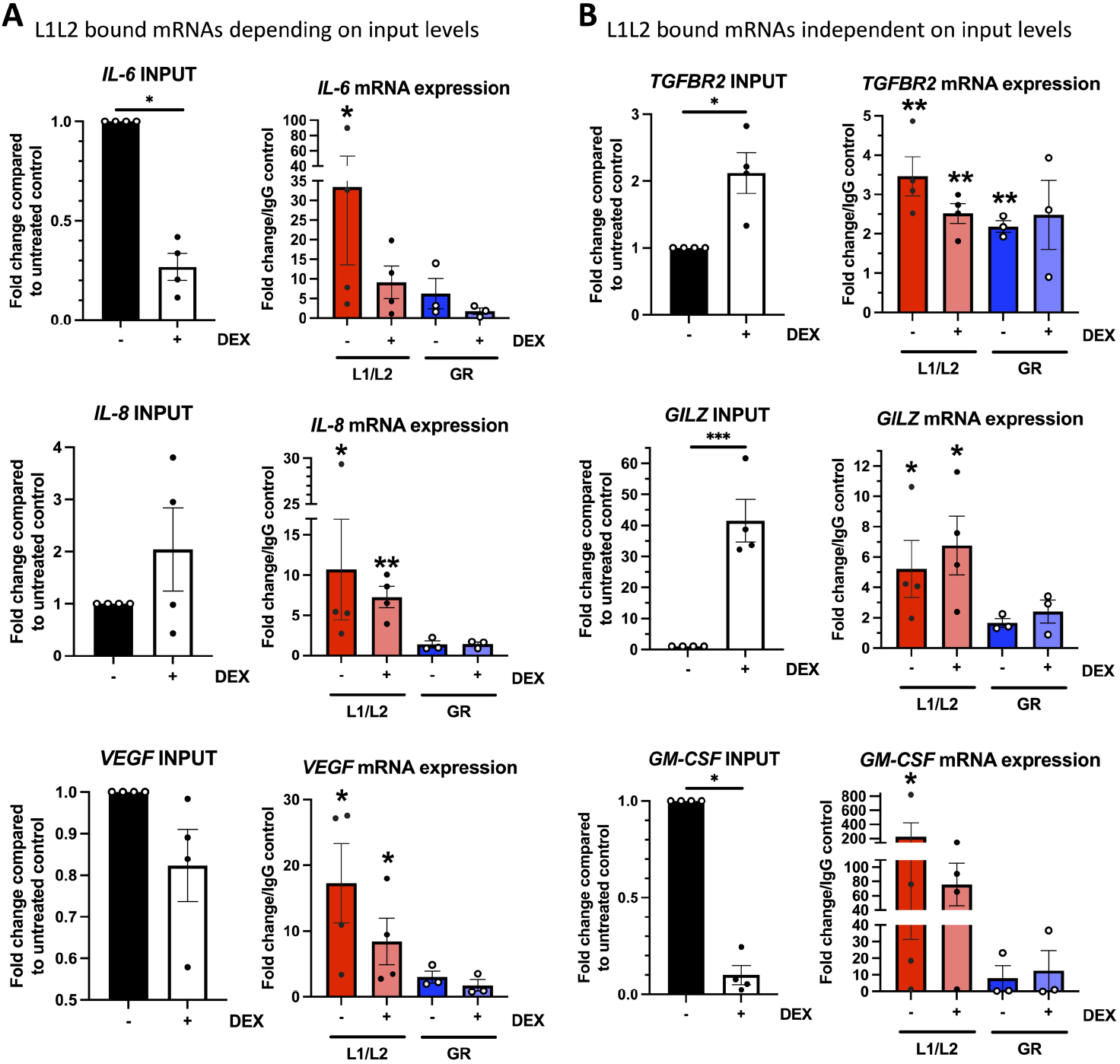
GC modulates ZFP36L1/L2 and GR binding to their mRNA targets. BEAS-2B cells were stimulated 10^−7^ M dexamethasone (+DEX) or vehicle (-DEX) for 24 hours and lysed for RNA immunoprecipitation. Bar graphs on the left represent the mean fold change of mRNA expression (±SEM) of the input samples. Bar graphs on the right represent mean fold change of mRNA expression (±SEM) over the respective IgG control (+ or -DEX). **A** depict mRNA candidates whose binding follow the pattern of expression triggered by GC stimulation: *IL6* (upper), *IL8* (middle) and *VEGF* (bottom) panels. **B** depict candidates whose biding does not reflect the changes triggered by GC and thus potentially modulated by GC: *TGFBR2* (upper), *GILZ* (middle) and *GM-CSF* (bottom). Summary data from three to four independent experiments. Statistical significance was assessed by multiple two-tailed t-tests on log transformed data comparing L1L2 immunoprecipitated mRNAs to their condition-matched (-/+ DEX) IgG control. p < 0.5 *, p < 0.01 ** p < 0.001 ***.

We also found that dexamethasone potentially modified the binding of ZFP36L1/L2 to some mRNAs. *TGFBR2* and *GILZ* mRNA levels were statistically increased by dexamethasone (Figure 2B upper and middle panels), while their binding to ZFP36L1/L2 remained the same and did not reflect this increase. *CSF2* (*GM-CSF* herein) mRNA was bound by ZFP36L1/L2; upon dexamethasone treatment *GM-CSF* mRNA levels were significantly decreased and showed no statistically significant binding to ZFP36L1/L2 (Figure 2B lower panels). Amongst all these transcripts, GR appeared to only bind *TGFBR2* mRNA in the absence of steroid (Figure 2B, upper panel blue bars) with a clear trend also in GC stimulated cells. Taken together, our data show that GC stimulation can modulate the binding of ZFP36L1/L2 and GR to their targets, and that the effects of GC transcriptionally may not reflect post-transcriptional processing of those mRNAs.

### Glucocorticoids exert transcriptional and post-transcriptional effects

Having established that GCs modulated both the levels and binding of ZFP36L1/L2 to their targets (Figures 1 and 2), we set out to determine their genome-wide role in the presence and absence of GCs. To that end, we used Frac-seq (subcellular <u>frac</u>tionation and RNA-<u>seq</u>uencing) (31). This allowed us to compare total mRNA, monosome-bound and polyribosome-bound mRNAs. Monosome-bound mRNAs may represent transcripts that have lower half-life, that require constant low-level translation into protein or that are undergoing elongation (32) while polyribosome (in our case more than five ribosomes) represent mRNAs engaged in translation. We chose to deplete both RBPs together, as they can regulate their own mRNAs considering that *ZFP36L1* and *ZFP36L2* mRNAs precipitated with ZFP36L1/L2 in bronchial epithelial cells (Supplemental Figure 2) and contain AREs. Human primary bronchial epithelial cells (BECs) were depleted of ZFP36L1 and ZFP36L2 employing siRNAs or siControl (scrambled siRNAs) as control, and both control and knockdown cells were treated with 10^−7^ M dexamethasone before being subjected to Frac-seq. Our polyribosome profiles showed that we could capture up to nine intact polyribosome peaks (Figure 3A and Supplemental Figure 3). Efficient knockdown of *ZFP36L1* and *ZFP36L2* was demonstrated at the total mRNA level by RT-qPCR (Figure 3B).

**Figure 3.**
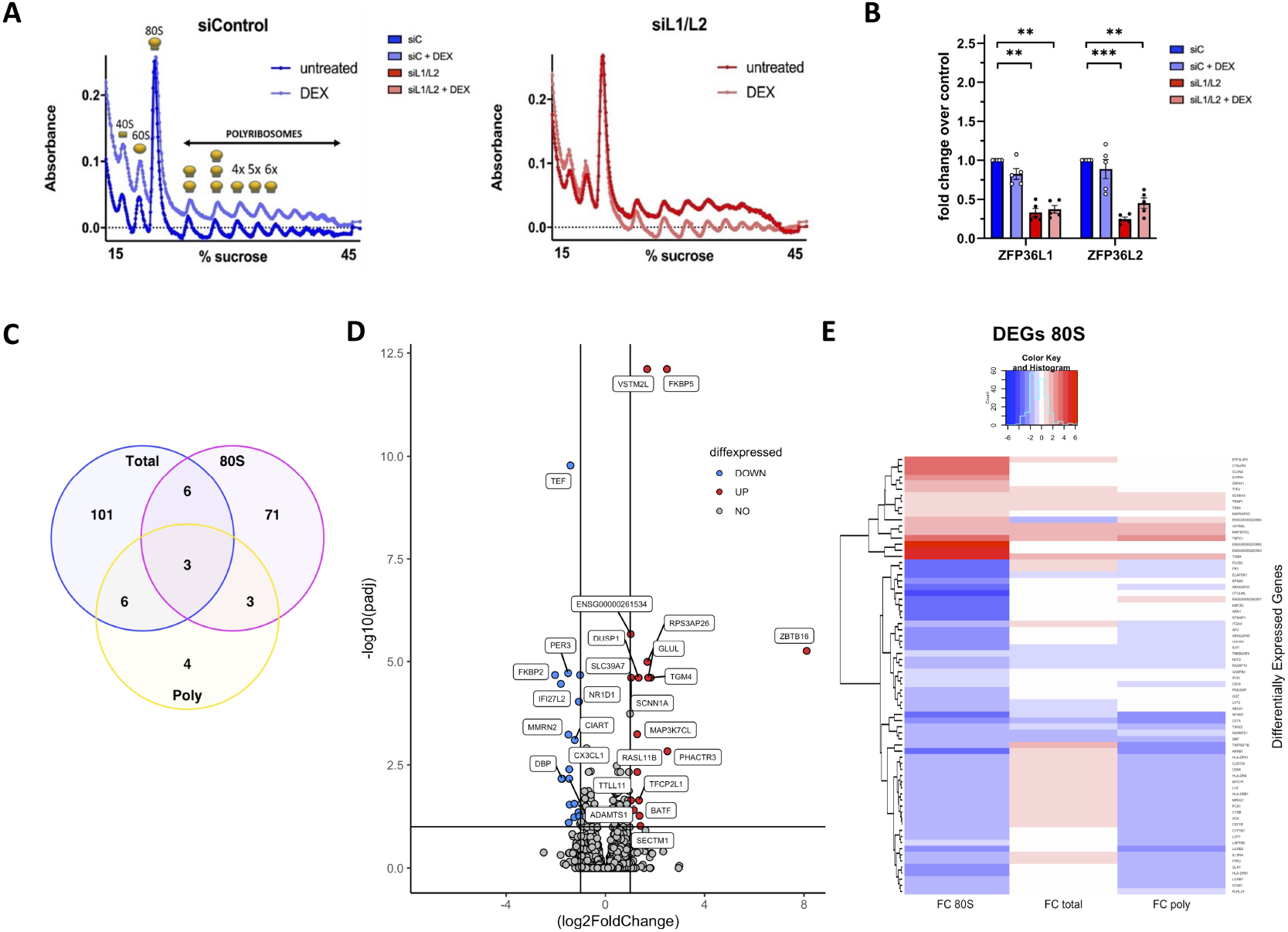
Frac-seq reveals that GC modulate transcription and 80S binding of distinct mRNAs. Primary BECs were transfected with siRNA control (siControl=blue lines) or siRNAs against ZFP36L1 and ZFP36L2 (siL1/L2=red lines) and stimulated with 10^−7^ M dexamethasone (DEX) or vehicle overnight. Cells were harvested and subjected to polyribosome fractionation and RNA sequencing. **A.** Representative polyribosome profiles from Frac-seq in primary BECs. Line graphs represent absorbance at 280 nm of mRNAs bound to ribosomes (Y-axis) across a density gradient (X-axis). Ribosomal subunits, units and polyribosome peaks (P1 onwards) are depicted. **B.** Total mRNA expression of *ZFP36L1* and *ZFP36L2* as quantified by RT-qPCR. Bar graphs represent fold changes (± SEM) of the five biological replicates sequenced, normalised to *GAPDH*. Values are relative to the untreated vehicle control. Statistical significance was assessed by two-tailed multiple t-tests on log transformed data. **C.** Venn diagram depicting the overlap of DEGs in response to dexamethasone in each fraction. Total = total mRNA, 80S = monosome-bound mRNA, Poly = polyribosome-bound mRNA. **D.** Volcano plot representing the DEGs (p-adjusted <0.1) in the 80S fraction in response to 10^−7^ M dexamethasone in control samples. Upregulated = red, downregulated = blue, grey = ns. **E.** Heatmap showing the fold change of DBGs in the 80S fraction (FC 80S, p-adjusted <0.1) in response to 10^−7^ M dexamethasone and their corresponding fold changes in total (FC total) and polyribosome (FC poly) fractions. p < 0.05 *, p < 0.01 ** p < 0.001 ***

Firstly, we set out to establish the transcriptional and post-transcriptional signatures exerted by GCs on primary BECs. We thus compared the effects of dexamethasone analyzing the samples from BECs transfected with siRNA control, in a fraction-dependent manner. The differentially expressed genes (DEGs, Supplemental Table 1) or differentially bound genes (DBGs, Supplemental Tables 2 and 3) were largely unique to each fraction (Figure 3C), with only three DEGs or DBGs (*FKBP5, DUSP1* and *ZBTB16*) in all three fractions upon dexamethasone treatment. These genes have previously been shown to be regulated by GCs (27). The Volcano plot on Figure 3D represents the transcriptional signature exerted by dexamethasone in these cells (which mapped to IL-4/IL-13/TGF-β/Wnt signalling when analysed using the Reactome database (33); these are all pathways known to be modulated by GCs at the level of transcription. GC did not appear to have a stark influence in polyribosome-bound mRNAs, with only 16 DBGs in that fraction.

Our previous Frac-seq work centred on comparing total vs polyribosome-bound mRNAs (31, 34), where we avoided isolating monosomes. Considering the addition of 80S to our analysis, we firstly validated that monosomes indeed contained a distinct population of transcripts. Previous work has shown that monosomes are able to translate mRNAs with low half-life and that monosome/polyribosome ratio serves as a measure of elongation rather than initiation (32). We firstly divided the baseMean counts from DESeq2 (in this case, the comparisons of dexamethasone effects in siControl transfected cells) to obtain an 80S/polyribosome ratio. We next filtered for genes that had more than 50 counts in monosomes and analysed the top and bottom 10% transcripts according to their monosome/polyribosome ratios. Supplemental Figure 4 shows that mRNAs with a low 80S/Poly ratio contained more than 99% proteincoding genes, while transcripts that showed a high 80S/polyribosome ratio had just 67.6% protein containing transcripts. This is consistent with previous reports showing that lncRNAs can bind to ribosomes where they may produce small peptides (35, 36) or be more prone to degradation (37). Thus, our 80S fraction consists of a distinct population of mRNAs and thus we analysed it as an independent fraction hereafter.

We discovered a high number of transcripts (83) regulated by dexamethasone in the 80S or monosomal fraction. 77% of monosome DBGs (64/83) were downregulated, and these corresponded to transcripts that were mostly increased in the total mRNA fraction (see changes from blue to red in the heatmap in Figure 3E).

### ZFP36L1/L2 exert distinct effects transcriptionally and post-transcriptionally in bronchial epithelial cells

We next analysed the changes exerted genome-wide by ZFP36L1/L2 depletion. We observed a large number of mRNAs modulated by ZFF36L1/L2 depletion (p-adjusted < 0.1) at the level of Total (662 DEGs) and polyribosome binding (133 DBGs) but very few at the level of 80S binding (9 DBGs), represented as volcano plots in Figure 4A. These data suggest that ZFP36L1/L2 exert distinct effects in a sub-cellular dependent manner, with a major role in cytoplasmic RNA levels and steady translation. Since ZFP36L1 and ZFP36L2 preferentially bind mRNAs containing AREs, we sought to explore if there was an enrichment for transcripts containing AREs upon ZFP36L1/L2 depletion, in a subcellular-dependent manner. We detected an enrichment of ARE-containing genes when compared to the expected frequency genome-wide at the total and polyribosome-binding levels (Figure 4B) based on the latest data available (38). We set out the expected frequency of ARE-containing genes at 25.55%, considering the ARE database (38) and 19,116 human protein coding genes (39). Considering only upregulated DEGs or DBGs, 35.8% DEGs in Total had AREs, while 33.3% and 32% were ARE-containing DBGs in the 80S and Poly fractions, respectively. The dysregulated ARE-containing genes upon ZFP36L1/L2 depletion also differed between fractions, highlighted in the Venn diagram in Figure 4C. Thus, ZFP36L1/L2 modulate ARE-containing gene expression altering their total RNA levels and also their binding to polyribosomes.

**Figure 4.**
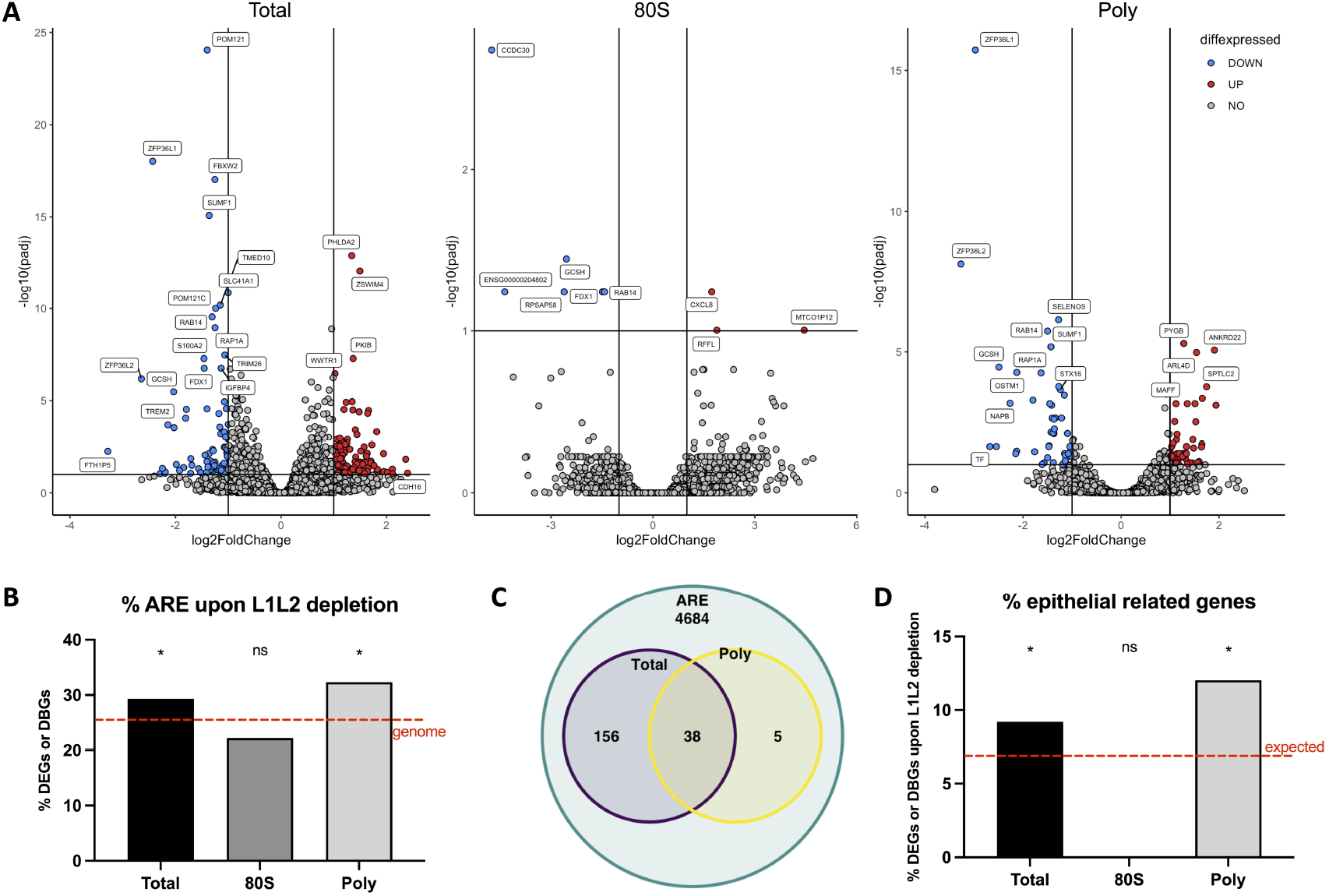
Frac-seq reveals that ZFP36L1/2 exert sub-cellular dependent effects. **A**. Volcano plots depicting DEGs (padj <0.1) in response to knock down of ZFP36L1/L2 in the total mRNA fraction (Total, left), the 80S fraction (80S, middle) and the polyribosome fraction (Poly, right). Upregulated = red, downregulated = blue, grey = ns. **B**. Percentage of dysregulated genes in Total, 80S and Poly that contain ARE elements. The red dotted line represents the ARE frequency in the human genome. Statistics were done using a one-tailed binomial test. **C**. Venn diagram highlighting the overlap of ARE-containing genes dysregulated from (A) in Total and Poly with the ARE database (38). **D**. Bar graph showing the percentage of transcripts that are modulated by ZFP36L1/L2 knock down that map to ‘epithelium’ in Gene Ontology per sub-cellular fraction. The red dotted line represents expected frequency genome-wide. Binomial two tailed test. p < 0.05 *, p < 0.01 ** p < 0.001 ***

ZFP36L1/L2 are best known for modulating immune-mediated processes (20, 21, 23, 25, 26, 40). We set out to determine if they play a role in the regulation of epithelial-related genes. To that end, we sought for the Gene Ontology term ‘epithelium’ in AmiGO2 (41–43) and we performed an enrichment analysis with the modulated candidates upon ZFP36L1/L2 down regulation, over the ‘expected’ frequency of these genes in the human genome (1,332 in 19,116 genes as per (39)). We detected an over-representation in total DEGs and polyribosome-bound DBGs (Figure 4D) suggesting that ZFP36L1/L2 can modulate the expression of epithelial-related genes such as ankyrin proteins, galectin like proteins and cadherins (for example *CDH16* and *PCDH1*).

### Glucocorticoids exert transcriptional and post-transcriptional effects in a ZFP36L1/L2-dependent manner

After analysing the genome-wide effects of GCs (Figure 3) and ZFP36L1/L2 depletion (Figure 4), we explored the possible interplay between ZFP36L1/L2 and GCs. GC binding to GR causes its nuclear translocation (9) and ZFP36L1 and ZFP36L2 can shuttle between nucleus and cytoplasm (44). Considering that GC appeared to regulate the binding of ZFP36L1/L2 to their targets (Figure 2B), we set out to determine if they modulated their subcellular localisation. BEAS-2B cells were stimulated with GC and immunofluorescence was undertaken at different time points, employing the dual ZFP36L1/L2 antibody. Figure 5A shows that ZFP36L1/L2 translocated to the nucleus in a time- and GC-dependent manner, suggesting a potential role for ZFP36L1/L2 in GC-mediated effects.

**Figure 5.**
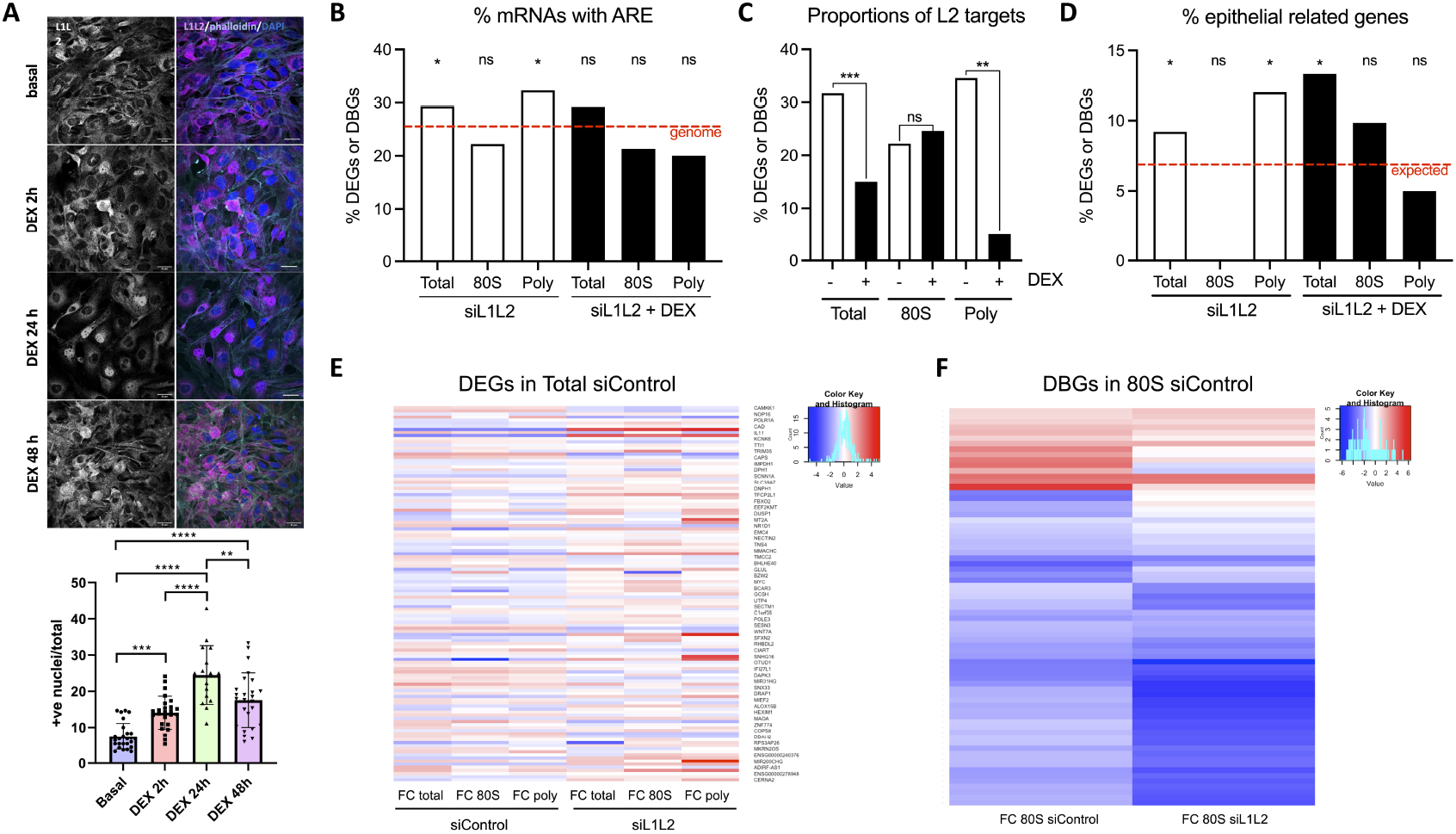
Frac-seq reveals that ZFP36L1/2 exert sub-cellular dependent effects and modulate GC action. **A**. Heatmap representing fold change in response to 10^−7^ M dexamethasone in all samples in each subcellular Upregulation by dexamethasone is depicted in red, while downregulation is depicted in blue. siControl = control-transfected; siL1L2: ZFP36L1/L2 knockdown. **B**. Confocal images of BEAS-2B cells after different exposure times to dexamethasone (DEX). Cells were stained with ZFP36L1/L2 antibody (L1L2, magenta), actin (phalloidin, cyan) and nucleus (DAPI, blue). The graph (mean ± SEM) shows the quantification of nuclear staining, pooling 3 different independent experiments. Each point in the graph represents a field with an average of 50 cells/field. A two-way ANOVA was used to compare the treatment groups. **C**: Bar graph showing the percentage of dysregulated genes in ZFP36L1/L2 knockdown cells in Total, 80S and Poly that contained ARE elements, in the absence (white, comparing siControl vs siL 1L2) or presence (black, comparing siControl + DEX vs siL1L2 + DEX) of dexamethasone. The red dotted line represents the ARE frequency in the human genome. Statistics were done using a one-tailed binomial test. The red dotted line represents the % ARE-containing genes genome-wide. **D**. Bar graph showing the effect of dexamethasone in the proportion of ZFP36L2 targets in Total, 80S and Poly in ZFP36L1/L2 depleted cells. Comparisons were done employing a two-sided Fisher’s exact test. **E**. Bar graph showing the effects of dexamethasone in ZFP36L1/L2 modulated genes that relate to epithelial cell function. The red dotted line represents the expected frequency genome-wide. Observed vs expected (genome-wide) frequencies comparisons were done employing a binomial two-tailed test. FC: fold change. p < 0.05 *, p < 0.01 ** p < 0.001 ***

We then analysed the presence of ZFP36L1/L2 targets upon dexamethasone stimulation and compared them with those of vehicle treated cells. The enrichment in ARE-containing DEGs and DBGs in Total and Poly upon ZFP36L1/L2 depletion (Figure 4B) was abolished by GC stimulation (Figure 5B). Although both Total (siControl and siL1L2) appeared the same, percentage-wise, the differences rely on the power of the analysis, i.e. 30% of 662 genes is more powerful statistically than 30% of 120 genes and this needs consideration when interpreting our data. Likewise, the ZFP36L2 targets from (40) were found significantly depleted in Total and Poly upon GC stimulation in ZFP36L1/L2 depleted samples (Figure 5C) while those in the 80S samples remained unmodulated. We also detected a depletion in the proportion of epithelial-related genes regulated by ZFP36L1/L2 (Figure 5D) in polyribosome-bound RNAs upon dexamethasone treatment, signifying the biological relevance of some of the mRNAs modulated by ZFP36L1/L2 in a GC-dependent manner.

To determine the dependence of GC action on ZFP36L1/L2 levels, we firstly analysed the overlap between the genes modulated by GC when comparing control to ZFP36L1/L2 depleted cells in a sub-cellular dependent manner (Supplemental Figure 5). We detected little overlap, with 31% of GC modulated genes in control cells also modulated upon ZFP36L1/L2 knockdown in Total, 7.2% in 80S and 43.8% in Poly. We then determined if the better-established transcriptional response to GCs was modulated by ZFP36L1/L2 levels in a sub-cellular dependent manner. We analysed the effects of GCs stimulation in Total (siControl Total comparing vehicle vs dexamethasone) and analysed the change exerted by GCs in those same DEGs in a sub-cellular and ZFP36L1/L2-dependent manner. Figure 5E shows a heatmap depicting these fold-changes in each fraction and transfection condition. As in Figure 3, we detected that GC differentially modulated mRNAs in a sub-cellular dependent manner (compare the first, second and third columns). Upon ZFP36L1/L2 depletion, GC exerted the opposite effects in myriad mRNAs (see change of colour from red to blue or blue to red between columns 1 vs 4, 5 or 6). Lastly, and considering the major role that we observed in GC modulating association to monosomes (Figure 3E), we analysed the DBGs to 80S upon GC stimulation from siControl cells with those in ZFP36L1/L2 depleted cells. Figure 5F shows that the widespread monosome association changes that GCs exerted showed a different pattern of modulation upon ZFP36L1/L2 knockdown. These data demonstrate that many of the sub-cellular dependent effects of GC (Figure 3) were modulated by ZFP36L1/L2 depletion in a sub-cellular dependent manner.

## Discussion

We present a new role for the RNA binding proteins ZFP36L1 and ZFP36L2 in the modulation of GC effects in bronchial epithelial cells, both transcriptionally and post-transcriptionally. GC modulated the levels of ZFP36L1 and ZFP36L2 in bronchial epithelium (Figure 1), their binding to specific transcripts (Figure 2) and increased their nuclear localisation (Figure 5A). Frac-seq revealed that gene expression regulation by GC mainly affects cytoplasmic levels and monosome association of distinct groups of mRNAs (Figure 3). Upon ZFP36L1/L2 depletion, the effects of GCs were modified genome-wide in a subcellular-dependent manner (Figures 5E and 5F). Our work uncovers that GCs are able to modify the proportions of both ARE-containing genes and known ZFP36L2 targets that are dependent on ZFP36L1/L2 levels (Figures 5B and 5C). Together, our data strongly suggest that GC and ZFP36L1/L2 are part of an intricate transcriptional and post-transcriptional network and uncover a new role for sub-cellular dependent effects of GCs on gene expression modulation via RBP regulation.

GCs are one of the most prescribed anti-inflammatory drugs worldwide. At the molecular level, GCs have been mainly described as transcriptional regulators, via binding to the GR and modulating the expression of genes that contain GREs in their promoters (8, 9, 27, 45). There is an increased appreciation of the role of GCs as post-transcriptional modulators, directly via GR as an RBP or via other RBPs such as ZFP36 or ZFP36L2 (12–14, 40, 46). There is also previous evidence that *ZFP36L1* and *ZFP36L2* mRNA or protein are upregulated in response to dexamethasone in some cell types (40, 47–49), but not bronchial epithelial cells so far. However, the effects of GCs, either transcriptional or post-transcriptional, have mostly being studied by assessing total mRNA levels. Considering the poor correlation between mRNA and protein levels (50–52) with polyribosome profiling showing a better proxy for protein production (53), we analysed the effects of GCs in different subcellular fractions, namely Total, 80S and Poly.

Our data revealed that GCs modulate cytoplasmic mRNA levels and association to monosomes, while polyribosome binding was only modified for a few genes (Figure 3). We performed further bioinformatic analysis seeking to determine potential mechanisms underlying the regulation of the 80S DBGs. We firstly assessed which DBGs had been previously described as GMD (16). We only found one hit, *NCF2*, whose binding to 80S was downregulated upon GC stimulation. Thus, GMD does not seem to drive the broad downregulation that we observed in monosome binding, although we cannot exclude the possibility that human BECs may have other GMD candidates. We also analysed if any DBGs in monosomes had been previously demonstrated to be directly bound by GR as per (14), however, we did not see any targets. Considering the discrepancy between the modulation of the levels of these genes by dexamethasone in the different fractions (Figure 3E), GC stimulation appeared to modulate monosome binding independently of total mRNA levels, suggesting that it has distinct transcriptional and post-transcriptional effects. Since the mammalian target of rapamycin (mTOR) is a major regulator of translation and there is some evidence of GC/GR crosstalk with mTOR (54, 55) we also analysed if DBGs in 80S had TOP (terminal oligopyrimidine) motifs (56), typically present in mRNAs regulated by mTOR. However, we did not see any TOP-containing mRNAs modulated by dexamethasone. Likewise, we detected no enrichment of IRES- (internal ribosome entry site) containing mRNAs in the 80S fraction over genome-wide levels (binomial test, p > 0.9999) (57).Considering that monosome association may serve as a readout for short lived RNAs (32), and that the majority of downregulated 80S DBGs appeared up-regulated in Total, 80S association may be used as a proxy for increased RNA decay. Based on the above, it is tempting to hypothesise that GCs modulate the stability of mRNA subsets via modulation of their 80S association.

Our data also allowed us to explore other roles for not only GCs but also for ZFP36L1/L2. ZFP36L1 and ZFP36L2 depletion appeared to modulate total cytoplasmic mRNA levels and association to polyribosomes, with little alterations in monosome association (Figure 4), in line with previous reports that these RBPs mainly modulate cytoplasmic RNA levels (stability) as well as translation (21, 22). We further interrogated the targets modulated by ZFP36L1/L2 and searched for epithelial related genes based on Gene Ontology and AmiGO2 (41–43) (Figures 4D and 5D). Our data suggest that ZFP36L1/L2 may modulate the expression of mRNAs that encode for epithelial-related functions. We understand that pathway analysis can be informative, but also have caveats and thus we also compared our datasets to previously published work. While we did not find many exact matches (e.g. SRM in Total DEGs, as in (23)), but genes upregulated upon *Zfp36l1/l2* deletion in mice thymus cells such as *Lrp1, Ptpn14, Notch1, Hes1* and *Lrp1* had family members upregulated in our dataset (*PTPN1, LRP8, HES4, HES5, NOTCH2NLC*). We also found *KLF2* upregulated upon ZFP36L1/L2 depletion as previous work on *Zfp36l2* over-expression on CD4+ T cells in mice (26). Moreover, we found family members of galectin like proteins, ankyrin proteins and cadherin proteins modulated in both our dataset and that of Vogel et al. (25). We also found that in ZFP36L1/L2 depleted samples, DEGs and DBGs were mapped to cell cycle pathways in line with previous findings (25, 58). Together, these data suggest a possible conservation of biological functions for ZFP36L1/L2 through modulation of tissue-specific gene family members in regulating cell integrity and structure.

GCs also appeared to modulate ZFP36L1/L2 binding to some transcripts (Figure 2). To our knowledge, our work is the first time this phenomenon has been assessed by RNA immunoprecipitation and that *GILZ* and *TGFBR2* have been identified as ZFP36L1/L2 targets. Our data also show how ZFP36L1/L2 can modulate their own encoding transcripts (Supplemental Figure 2). GR also showed significant binding to *ZFP36L1* and *ZFP36L2* mRNA in the absence of dexamethasone (Supplemental Figure 2) and a trend towards association upon dexamethasone stimulation. GC treatment caused a similar number of DEGs and DBGs in control and ZFP36L1/L2 depleted samples; however, the overlap between them was little (Supplemental Figure 5), particularly in the Total and 80S fractions. Moreover, our data strongly suggest that GC modulated the ZFP36L1/L2-dependent ARE-containing transcripts and previously published ZFP36L2 targets genome-wide (Figures 5B and 5C) in a sub-cellular dependent manner.

Together, our data suggest that ZFP36L1/L2 depletion impacted GC-mediated responses in a subcellular dependent manner, where nearly 70% of DEGs by GC in Total and over 90% of DBGs in 80S were no longer modulated by GC upon ZFP36L1/L2 depletion (Supplementary Figure 5). Thus, the most studied fraction to understand GC effects, Total, overlooks the effects that GC causes in other subcellular compartments, and these changes are dependent on ZFP36L1/L2 levels (Figure 5E). Concomitantly, we observed a shift towards nuclear localisation for ZFP36L1/L2 (Figure 5A) suggesting that GC modulate gene expression not only via effects on GREs or decay, but also via subcellular compartmentalisation of these RBPs.

In summary, our work demonstrates that GC exert subcellular dependent effects, with monosome binding as a novel subcellular compartment modulated by GC. ZFP36L1/L2 levels are able to modulate the genome-wide response of GCs, both transcriptional and post-transcriptionally. Improving our understanding of these RBPs may lead to an improvement in GC effectiveness, GC-response biomarker discovery and/or potential tailoring of GC therapy. Our data show the relevance and feasibility of employing sub-cellular fractionation to fully understand the role of RBPs in genome-wide regulation of mRNA expression, and to reveal novel pathways and effects of stimuli that are ‘classical’ transcriptional mediators.

## Materials and Methods

### Cell culture

BEAS-2B cells (American Type Culture Collection; ATCC) were grown in Roswell Park Memorial Institute (RPMI) (Invitrogen) medium with GlutaMAX supplemented with 10% fetal bovine serum (FBS) (Sigma-Aldrich). Primary bronchial epithelial cells (BECs) we obtained under ethical approval (REC Number 06/Q0505/12) and grown in airway epithelial cell medium (Promo Cell) supplemented bovine pituitary extract, epidermal growth factor, insulin, hydrocortisone, epinephrine, triiodo-L-thyronine, transferrin, and retinoic acid on collagen-coated plates. All cells were incubated at 37 °C in 5% CO_2_ and were passaged when 80% confluent. Prior to stimulation, BEAS-2B cells were serum starved in 2% FBS RPMI overnight and BECs in complete medium minus hydrocortisone. Unless otherwise stated, 10^−7^ M Dexamethasone was used for GC treatment.

### siRNA transfections

siRNAs (Silencer Select, ThermoFisher Scientific) were used at a final concentration of 25 nM. Interferin (Polyplus) was used as a transfection reagent following manufacturer’s instructions. Cells were incubated in the transfection mix for 4 hours before being replenished in 2% FBS RPMI or BEC medium without hydrocortisone in preparation for stimulation with 10^−7^ M dexamethasone 24h later.

### Reverse transcription and quantitative PCR (RT-qPCR)

RNA was extracted using TRIzol (Invitrogen) or TRIzol LS (Sigma-Aldrich) and reverse transcribed employing RNAse H Minus Reverse Transcriptase, Ribolock and random hexamers (ThermoFisher Scientific). qPCR was performed using NEB Luna buffer (New England Biolabs) and TaqMan assays (ThermoFisher Scientific). Gene expression was normalised against *GAPDH* (Primer Design).

### Western blot analysis

Cells were lysed on ice in protein lysis buffer (0.5% NP40, 20mM Tris HCl pH 7.5, 100mM KCl, 10mM MgCl_2_) supplemented with fresh protease inhibitor cocktail (Cell Signalling). Lysate was loaded and resolved by SDS-PAGE on a 12% Bis-Acrylamide gel and electrophoresis was carried out at using NuPAGE MES SDS running buffer (ThermoFisher Scientific). The separated proteins were transferred onto a nitrocellulose membrane (BioRad Laboratories) at 40V for 90 minutes using a wet transfer system and NuPAGE transfer buffer (ThermoFisher Scientific) containing 20% methanol. The membrane was incubated in blocking buffer (Amersham Biosciences) for 60 minutes at room temperature and in primary antibody (BRF1/2: CST#2119, GR: CST#3660, GAPDH: Santa Cruz sc32233) overnight at 4°C. This was followed by incubation with the appropriate anti-rabbit or anti-mouse IgG antibody conjugated to horseradish peroxidase (Dako) for 60 minutes at room temperature. All washes and antibody dilutions were performed in TBS-0.05% Tween. Immunoreactive proteins were detected using the ECL immunodetection system (ThermoFisher Scientific) and visualised using the ChemiDoc MP Imaging System (BioRad) and ImageLab software for analysis.

### Immunofluorescence

Cells on coated coverslips were plated, and 24 hours later there were washed with PBS, fixed with 4% PFA in PBS for 10 min and permeabilised with 0.2% TritonX-100 for 10 min. Cells were incubated with primary antibodies overnight and appropriate secondary antibodies and Phalloidin conjugated to Alexafluor 568 or 633 for 1 hour. Cells were mounted onto slides using FluorSave ® (Millipore). Confocal microscopy was performed using a Nikon A1R inverted confocal laser scanning microscope with a 60x oil objective and laser excitation wavelength of 488 nm (for GFP or Alexafluor-488), 561 nm (for Alexafluor-568) and 633 nm (for Alexafluor-633 and cy5). Images were exported from the Nikon Elements software (Nikon) for further analysis in ImageJ software (U. S. National Institutes of Health, Bethesda, MD, USA).

### RNA immunoprecipitation (RIP)

RIP was performed with 1 μg antibody. Antibodies used were: BRF1/2 (Cell Signaling Technology #2119), GR (Cell Signaling Technology #3660) and Rabbit IgG (Merck, #03198) per 1.5 million cells. Cell lysates were prepared from BEAS-2B cells treated with or without dexamethasone and lysed on ice in RIP lysis buffer (100 mM KCl, 25 mM Tris pH 7.4, 5 mM EDTA, 0.5 mM DTT, 0.5% NP40, 100 U/mL Ribolock RNase inhibitor (ThermoFisher Scientific), 1X protease inhibitor cocktail (Cell Signaling Technology). 10% of lysate volume was kept as the input. Dynabeads Protein A magnetic beads (ThermoFisher Scientific) were utilised for antibody binding and the antibody-bead complex was incubated on a rotating shaker for 10 minutes at room temperature. Cell lysates were immunoprecipitated with the antibody-bead complex on a rotating shaker overnight at 4°C with the addition of 100 U/mL Ribolock (ThermoFisher Scientific). RNA was extracted from beads by using TRIzol (ThermoFisher Scientific) and RNA presence was measured by RT-qPCR.

### Co-Immunoprecipitation

Following stimulation with dexamethasone overnight, BEAS-2B cells were washed with ice-cold PBS and lysed with 1 X Cell Lysis Buffer (Cell Signaling Technologies) supplemented with fresh protease inhibitor cocktail (Cell Signaling Technologies). Cells were left on ice for 5 mins, lysates passed through a 23G needle three times and then spun at 14,000 x g for 10 mins at 4°C. Approximately 10% of total lysate was saved for use as loading controls and the remainder was precleared with 20 μL of Protein A Dynabeads (ThermoFisher Scientific). Precleared lysates were immunoprecipitated with BRF1/2 antibody (#2119, Cell Signaling Technology), Glucocorticoid Receptor antibody (#3660, Cell Signalling Technology) or control IgG antibody (#03-198, Merck Millipore) and incubated for 1 h at room temperature whilst rotating. 50 μl of Protein A Dynabeads were added to each immunoprecipitation sample and incubated for 20 mins at room temperature. Samples were separated using a magnetic rack and washed three times with ice cold lysis buffer. Protein was eluted in 30 μL of protein loading buffer and heated at 95°C for 5 min. Eluted proteins were separated by SDS-PAGE, and analyzed by western blotting as described above.

### Polyribosome profiling

Polyribosome profiling was performed as described previously (31). Briefly BECs were transfected with siRNAs and stimulated with 10^−7^ M dexamethasone (Sigma Aldrich). 500 μg/ml cyclohexamide (Sigma-Aldrich) was added for 10 minutes prior to lysing. Supernatants were carefully removed and kept for cytokine analysis. BECs were lysed in polyribosome lysis buffer (0.5% NP40, 20mM Tris HCl pH 7.5, 100mM KCl, 10mM MgCl2, 1X protease inhibitor cocktail (Cell Signalling, 5871S), 100 μg/mL cycloheximide) on ice for 7 minutes and passed through a 23G needle three times. Cytoplasmic extracts were spun at 8000 rpm at 4°C for 7 minutes. 10% of cell lysate was saved for cytoplasmic (Total) RNA extraction; the rest was carefully loaded onto 15-45% sucrose gradients and then ultracentrifuged using TH-641 swinging buckets in a Hitachi ultracentrifuge at 40,000 rpm at 4°C for 1 hour 25 minutes. Preparation of the gradients, reading of polyribosome profiles, and extraction of individual polyribosome peaks were all done employing a Gradient Station (BioComp Instruments, Fredericton, NB, Canada) equipped with a Triax. Polyribosome profiles were generated reading the absorbance at 260 nm of spun gradients immediately after ultracentrifugation. Fractions corresponding to each ribosomal peak were fractionated and extracted using TRIzol LS (Sigma-Aldrich) following manufacturer’s instructions. The RNA from fractions P5 (6 ribosomes) onwards was pooled to constitute the Polyribosome fraction and was sequenced along with the total RNA and 80S monosomal fractions.

### Library preparation and RNA-sequencing

RNA was digested on-column using DNase (Qiagen) prior to library preparation using the NEBNext Ultra II Directional RNA Library Prep Kit for Illumina (New England Biolabs) with poly(A) mRNA selection and dual-index primers. 10 ng RNA was used as a starting input. Libraries were sequenced on a NovaSeq6000 (Illumina) utilising 150 bp sequencing to a depth of 40 M paired-end reads.

### RNA-seq analysis

Cancer Genomics Cloud (CGC) was used for data analysis. Sequencing data quality control was carried out with FastQC. Trimmomatic *(59)* was used to remove adapter sequences. Salmon index and Salmon quant from salmon (60) were used for alignment to the genome (v37). R studio was used to perform DESeq2 (61) for analysis of differential gene expression and all downstream data visualisation. Data was generated from five biological replicates. Gene levels were expressed as log_2_FoldChange and p-adjusted <0.1 (*p-adj*) was considered as significantly regulated. Datasets are available at GEO at the Reviewers’ request.

### Statistical analysis

GraphPad Prism and R studio software was used for the generation of graphs and analysis of data. All packages are available at CRAN and the code employed at GitHub https://github.com/rynne94?tab=repositories. Data is shown as mean ± standard error of the mean. Data was tested for normality using a Shapiro-Wilk normality test. Data was log transformed for normalisation and statistical analysis was performed using t-tests on the log transformed data (unless otherwise stated) to determine statistical significance between conditions. Where stated, a paired ratio test was performed. Enrichment analysis was performed using binomial tests. In all cases, p < 0.05 *, p < 0.01 **, p < 0.001 ***. RNA-seq analysis was evaluated using the DESeq2 package for differential gene expression, using an adjusted p-value of *p-adj* < 0.1.

## Acknowledgements

The authors thank Prof Michael H. Malim for his critical and helpful input in the manuscript. This work was funded by King’s Health Partners (Challenge Fund to RTMN), the Rosetrees Trust (JS15 / M726 to RTMN), the Wellcome Trust (213984/Z/18/Z) to RTMN and Asthma UK grant to fund JR PhD studentship and. For the purpose of open access, the author has applied a CC BY public copyright licence to any Author Accepted Manuscript version arising from this submission. The funders had no role in study design, data collection and analysis, decision to publish, or preparation of the manuscript.

## References

1. Benard-Laribiere A, Pariente A, Pambrun E, Begaud B, Fardet L, Noize P. Prevalence and prescription patterns of oral glucocorticoids in adults: a retrospective cross-sectional and cohort analysis in France. BMJ Open. 2017;7(7):e015905.

2. Laugesen K, Jorgensen JOL, Petersen I, Sorensen HT. Fifteen-year nationwide trends in systemic glucocorticoid drug use in Denmark. Eur J Endocrinol. 2019;181(3):267–73.

3. van Staa TP, Leufkens HG, Abenhaim L, Begaud B, Zhang B, Cooper C. Use of oral corticosteroids in the United Kingdom. QJM. 2000;93(2):105–11.

4. Volmer T, Effenberger T, Trautner C, Buhl R. Consequences of longterm oral corticosteroid therapy and its side-effects in severe asthma in adults: a focused review of the impact data in the literature. Eur Respir J. 2018;52(4).

5. De Bosscher K, Haegeman G. Minireview: latest perspectives on antiinflammatory actions of glucocorticoids. Mol Endocrinol. 2009;23(3):281–91.

6. Zhang T, Haws P, Wu Q. Multiple variable first exons: a mechanism for cell-and tissue-specific gene regulation. Genome Res. 2004;14(1):79–89.

7. Chen W, Dang T, Blind RD, Wang Z, Cavasotto CN, Hittelman AB, et al. Glucocorticoid receptor phosphorylation differentially affects target gene expression. Mol Endocrinol. 2008;22(8):1754–66.

8. Pratt WB, Morishima Y, Murphy M, Harrell M. Chaperoning of glucocorticoid receptors. Handb Exp Pharmacol. 2006(172):111–38.

9. Oakley RH, Cidlowski JA. The biology of the glucocorticoid receptor: new signaling mechanisms in health and disease. J Allergy Clin Immunol. 2013;132(5):1033–44.

10. Chinenov Y, Rogatsky I. Glucocorticoids and the innate immune system: crosstalk with the toll-like receptor signaling network. Mol Cell Endocrinol. 2007;275(1-2):30–42.

11. Hebbar PB, Archer TK. Chromatin remodeling by nuclear receptors. Chromosoma. 2003;111(8):495–504.

12. Ishmael FT, Fang X, Galdiero MR, Atasoy U, Rigby WF, Gorospe M, et al. Role of the RNA-binding protein tristetraprolin in glucocorticoid-mediated gene regulation. J Immunol. 2008;180(12):8342–53.

13. Stellato C. Post-transcriptional and nongenomic effects of glucocorticoids. Proc Am Thorac Soc. 2004;1(3):255–63.

14. Ishmael FT, Fang X, Houser KR, Pearce K, Abdelmohsen K, Zhan M, et al. The human glucocorticoid receptor as an RNA-binding protein: global analysis of glucocorticoid receptor-associated transcripts and identification of a target RNA motif. J Immunol. 2011;186(2):1189–98.

15. Dhawan L, Liu B, Blaxall BC, Taubman MB. A novel role for the glucocorticoid receptor in the regulation of monocyte chemoattractant protein-1 mRNA stability. J Biol Chem. 2007;282(14):10146–52.

16. Cho H, Park OH, Park J, Ryu I, Kim J, Ko J, et al. Glucocorticoid receptor interacts with PNRC2 in a ligand-dependent manner to recruit UPF1 for rapid mRNA degradation. Proc Natl Acad Sci U S A. 2015;112(13):E1540–9.

17. Park OH, Park J, Yu M, An HT, Ko J, Kim YK. Identification and molecular characterization of cellular factors required for glucocorticoid receptor-mediated mRNA decay. Genes Dev. 2016;30(18):2093–105.

18. Blackshear PJ. Tristetraprolin and other CCCH tandem zinc-finger proteins in the regulation of mRNA turnover. Biochem Soc Trans. 2002;30(Pt 6):945–52.

19. Cassandri M, Smirnov A, Novelli F, Pitolli C, Agostini M, Malewicz M, et al. Zinc-finger proteins in health and disease. Cell Death Discov. 2017;3:17071.

20. Lai WS, Carballo E, Strum JR, Kennington EA, Phillips RS, Blackshear PJ. Evidence that tristetraprolin binds to AU-rich elements and promotes the deadenylation and destabilization of tumor necrosis factor alpha mRNA. Mol Cell Biol. 1999;19(6):4311–23.

21. Salerno F, Engels S, van den Biggelaar M, van Alphen FPJ, Guislain A, Zhao W, et al. Translational repression of pre-formed cytokine-encoding mRNA prevents chronic activation of memory T cells. Nat Immunol. 2018;19(8):828–37.

22. Adachi S, Homoto M, Tanaka R, Hioki Y, Murakami H, Suga H, et al. ZFP36L1 and ZFP36L2 control LDLR mRNA stability via the ERK-RSK pathway. Nucleic Acids Res. 2014;42(15):10037–49.

23. Hodson DJ, Janas ML, Galloway A, Bell SE, Andrews S, Li CM, et al. Deletion of the RNA-binding proteins ZFP36L1 and ZFP36L2 leads to perturbed thymic development and T lymphoblastic leukemia. Nat Immunol. 2010;11(8):717–24.

24. Lin RJ, Huang CH, Liu PC, Lin IC, Huang YL, Chen AY, et al. Zinc finger protein ZFP36L1 inhibits influenza A virus through translational repression by targeting HA, M and NS RNA transcripts. Nucleic Acids Res. 2020;48(13):7371–84.

25. Vogel KU, Bell LS, Galloway A, Ahlfors H, Turner M. The RNA-Binding Proteins Zfp36l1 and Zfp36l2 Enforce the Thymic beta-Selection Checkpoint by Limiting DNA Damage Response Signaling and Cell Cycle Progression. J Immunol. 2016;197(7):2673–85.

26. Makita S, Takatori H, Iwata A, Tanaka S, Furuta S, Ikeda K, et al. RNA-Binding Protein ZFP36L2 Downregulates Helios Expression and Suppresses the Function of Regulatory T Cells. Front Immunol. 2020;11:1291.

27. Mostafa MM, Rider CF, Shah S, Traves SL, Gordon PMK, Miller-Larsson A, et al. Glucocorticoid-driven transcriptomes in human airway epithelial cells: commonalities, differences and functional insight from cell lines and primary cells. BMC Med Genomics. 2019;12(1):29.

28. Eddleston J, Herschbach J, Wagelie-Steffen AL, Christiansen SC, Zuraw BL. The anti-inflammatory effect of glucocorticoids is mediated by glucocorticoid-induced leucine zipper in epithelial cells. J Allergy Clin Immunol. 2007;119(1):115–22.

29. Silva CM, Powell-Oliver FE, Jewell CM, Sar M, Allgood VE, Cidlowski JA. Regulation of the human glucocorticoid receptor by long-term and chronic treatment with glucocorticoid. Steroids. 1994;59(7):436–42.

30. Vaughan MB, Ramirez RD, Wright WE, Minna JD, Shay JW. A threedimensional model of differentiation of immortalized human bronchial epithelial cells. Differentiation. 2006;74(4):141–8.

31. Sterne-Weiler T, Martinez-Nunez RT, Howard JM, Cvitovik I, Katzman S, Tariq MA, et al. Frac-seq reveals isoform-specific recruitment to polyribosomes. Genome Res. 2013;23(10):1615–23.

32. Heyer EE, Moore MJ. Redefining the Translational Status of 80S Monosomes. Cell. 2016;164(4):757–69.

33. Gillespie M, Jassal B, Stephan R, Milacic M, Rothfels K, Senff-Ribeiro A, et al. The reactome pathway knowledgebase 2022. Nucleic Acids Res. 2022;50(D1):D687–D92.

34. Martinez-Nunez RT, Rupani H, Plate M, Niranjan M, Chambers RC, Howarth PH, et al. Genome-Wide Posttranscriptional Dysregulation by MicroRNAs in Human Asthma as Revealed by Frac-seq. J Immunol. 2018;201(1):251–63.

35. Ji Z, Song R, Regev A, Struhl K. Many lncRNAs, 5′UTRs, and pseudogenes are translated and some are likely to express functional proteins. Elife. 2015;4:e08890.

36. Minati L, Firrito C, Del Piano A, Peretti A, Sidoli S, Peroni D, et al. One-shot analysis of translated mammalian lncRNAs with AHARIBO. Elife. 2021;10.

37. Carlevaro-Fita J, Rahim A, Guigo R, Vardy LA, Johnson R. Cytoplasmic long noncoding RNAs are frequently bound to and degraded at ribosomes in human cells. RNA. 2016;22(6):867–82.

38. Bakheet T, Hitti E, Khabar KSA. ARED-Plus: an updated and expanded database of AU-rich element-containing mRNAs and pre-mRNAs. Nucleic Acids Res. 2018;46(D1):D218–D20.

39. Piovesan A, Antonaros F, Vitale L, Strippoli P, Pelleri MC, Caracausi M. Human protein-coding genes and gene feature statistics in 2019. BMC Res Notes. 2019;12(1):315.

40. Zhang L, Prak L, Rayon-Estrada V, Thiru P, Flygare J, Lim B, et al. ZFP36L2 is required for self-renewal of early burst-forming unit erythroid progenitors. Nature. 2013;499(7456):92–6.

41. Ashburner M, Ball CA, Blake JA, Botstein D, Butler H, Cherry JM, et al. Gene ontology: tool for the unification of biology. The Gene Ontology Consortium. Nat Genet. 2000;25(1):25–9.

42. Gene Ontology C. The Gene Ontology resource: enriching a GOld mine. Nucleic Acids Res. 2021;49(D1):D325–D34.

43. Carbon S, Ireland A, Mungall CJ, Shu S, Marshall B, Lewis S, et al. AmiGO: online access to ontology and annotation data. Bioinformatics. 2009;25(2):288–9.

44. Phillips RS, Ramos SB, Blackshear PJ. Members of the tristetraprolin family of tandem CCCH zinc finger proteins exhibit CRM1-dependent nucleocytoplasmic shuttling. J Biol Chem. 2002;277(13):11606–13.

45. Timmermans S, Souffriau J, Libert C. A General Introduction to Glucocorticoid Biology. Front Immunol. 2019;10:1545.

46. Pemmari A, Leppanen T, Hamalainen M, Moilanen T, Vuolteenaho K, Moilanen E. Widespread regulation of gene expression by glucocorticoids in chondrocytes from patients with osteoarthritis as determined by RNA-Seq. Arthritis Res Ther. 2020;22(1):271.

47. Pennanen PT, Sarvilinna NS, Purmonen SR, Ylikomi TJ. Changes in protein tyrosine phosphatase type IVA member 1 and zinc finger protein 36 C3H type-like 1 expression demonstrate altered estrogen and progestin effect in medroxyprogesterone acetate-resistant and estrogen-independent breast cancer cell models. Steroids. 2009;74(4-5):404–9.

48. Sjogren SE, Siva K, Soneji S, George AJ, Winkler M, Jaako P, et al. Glucocorticoids improve erythroid progenitor maintenance and dampen Trp53 response in a mouse model of Diamond-Blackfan anaemia. Br J Haematol. 2015;171(4):517–29.

49. Hacker C, Valchanova R, Adams S, Munz B. ZFP36L1 is regulated by growth factors and cytokines in keratinocytes and influences their VEGF production. Growth Factors. 2010;28(3):178–90.

50. Brion C, Lutz SM, Albert FW. Simultaneous quantification of mRNA and protein in single cells reveals post-transcriptional effects of genetic variation. Elife. 2020;9.

51. Koussounadis A, Langdon SP, Um IH, Harrison DJ, Smith VA. Relationship between differentially expressed mRNA and mRNA-protein correlations in a xenograft model system. Sci Rep. 2015;5:10775.

52. Gygi SP, Rochon Y, Franza BR, Aebersold R. Correlation between protein and mRNA abundance in yeast. Mol Cell Biol. 1999;19(3):1720–30.

53. Wang T, Cui Y, Jin J, Guo J, Wang G, Yin X, et al. Translating mRNAs strongly correlate to proteins in a multivariate manner and their translation ratios are phenotype specific. Nucleic Acids Res. 2013;41(9):4743–54.

54. Lesovaya E, Agarwal S, Readhead B, Vinokour E, Baida G, Bhalla P, et al. Rapamycin Modulates Glucocorticoid Receptor Function, Blocks Atrophogene REDD1, and Protects Skin from Steroid Atrophy. J Invest Dermatol. 2018;138(9):1935–44.

55. Polman JA, Hunter RG, Speksnijder N, van den Oever JM, Korobko OB, McEwen BS, et al. Glucocorticoids modulate the mTOR pathway in the hippocampus: differential effects depending on stress history. Endocrinology. 2012;153(9):4317–27.

56. Cockman E, Anderson P, Ivanov P. TOP mRNPs: Molecular Mechanisms and Principles of Regulation. Biomolecules. 2020;10(7).

57. Zhao J, Li Y, Wang C, Zhang H, Zhang H, Jiang B, et al. IRESbase: A Comprehensive Database of Experimentally Validated Internal Ribosome Entry Sites. Genomics Proteomics Bioinformatics. 2020;18(2):129–39.

58. Galloway A, Saveliev A, Lukasiak S, Hodson DJ, Bolland D, Balmanno K, et al. RNA-binding proteins ZFP36L1 and ZFP36L2 promote cell quiescence. Science. 2016;352(6284):453–9.

59. Bolger AM, Lohse M, Usadel B. Trimmomatic: a flexible trimmer for Illumina sequence data. Bioinformatics. 2014;30(15):2114–20.

60. Patro R, Duggal G, Love MI, Irizarry RA, Kingsford C. Salmon provides fast and bias-aware quantification of transcript expression. Nat Methods. 2017;14(4):417–9.

61. Love MI, Huber W, Anders S. Moderated estimation of fold change and dispersion for RNA-seq data with DESeq2. Genome Biol. 2014;15(12):550.

