## Supplemental Figures for "The RNA binding proteins ZFP36L1 and ZFP36L2 modulate transcriptional and post-transcriptional genome-wide effects of glucocorticoids"

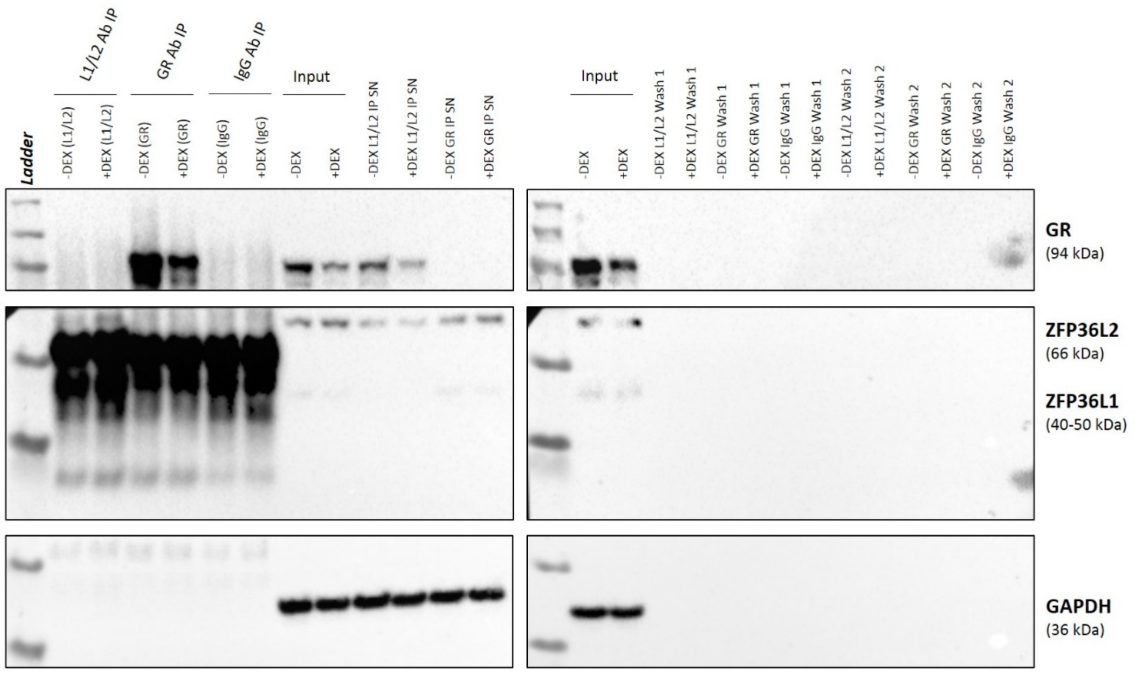


**Supplemental Figure 1. Immunoprecipitation analysis of L1/L2 protein interactions.** BEAS-2B cells were stimulated with $10^{-7}$ M dexamethasone (+DEX) or vehicle (- DEX), for 24 hours and lysed for immunoprecipitation. Lysates were precipitated with antibodies against ZFP36L1/L2 (L1/L2), GR, or an IgG control antibody. Lysates were subjected to SDS-PAGE and western blot analysis with antibodies against ZFP36L1, ZFP36L2 and GR. Total protein was used as the input control. Data represents one biological replicate. SN = supernatant, IP = immunoprecipitation.

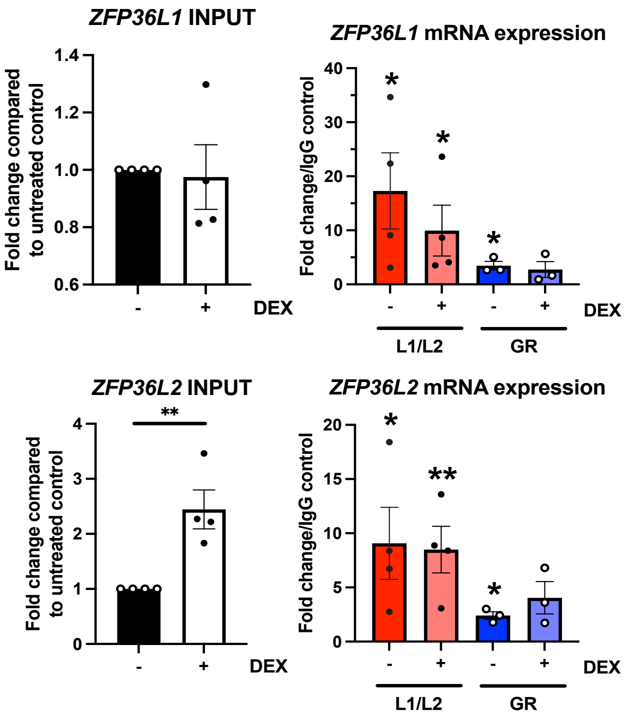


**Supplemental Figure 2. ZFP36L1 and ZFP36L2 mRNAs are bound by ZFP36L1/L2 proteins.** BEAS-2B cells were stimulated with 10-7 M dexamethasone (+DEX) or vehicle (-DEX) for 24 hours and lysed for RNA immunoprecipitation. ZFP36L1/ZFP36L2 and GR were immunoprecipitated with their associated RNAs to identify bound transcripts and detected by quantitative PCR for mRNA expression of ZFP36L1 (upper) and ZFP36L2 (lower). Bar graphs on the left represent the fold change (± SEM) of the input samples to identify changes in response to dexamethasone treatment irrespective of RBP binding capacity. Bar graphs on the right represent mean fold change (± SEM) over the respective IgG control of three to four independent experiments. Statistical significance was assessed by multiple two-tailed t-tests on log transformed data comparing DEX treatment (black asterisks). In all cases $p < 0.05$ *, $p < 0.01$ **.

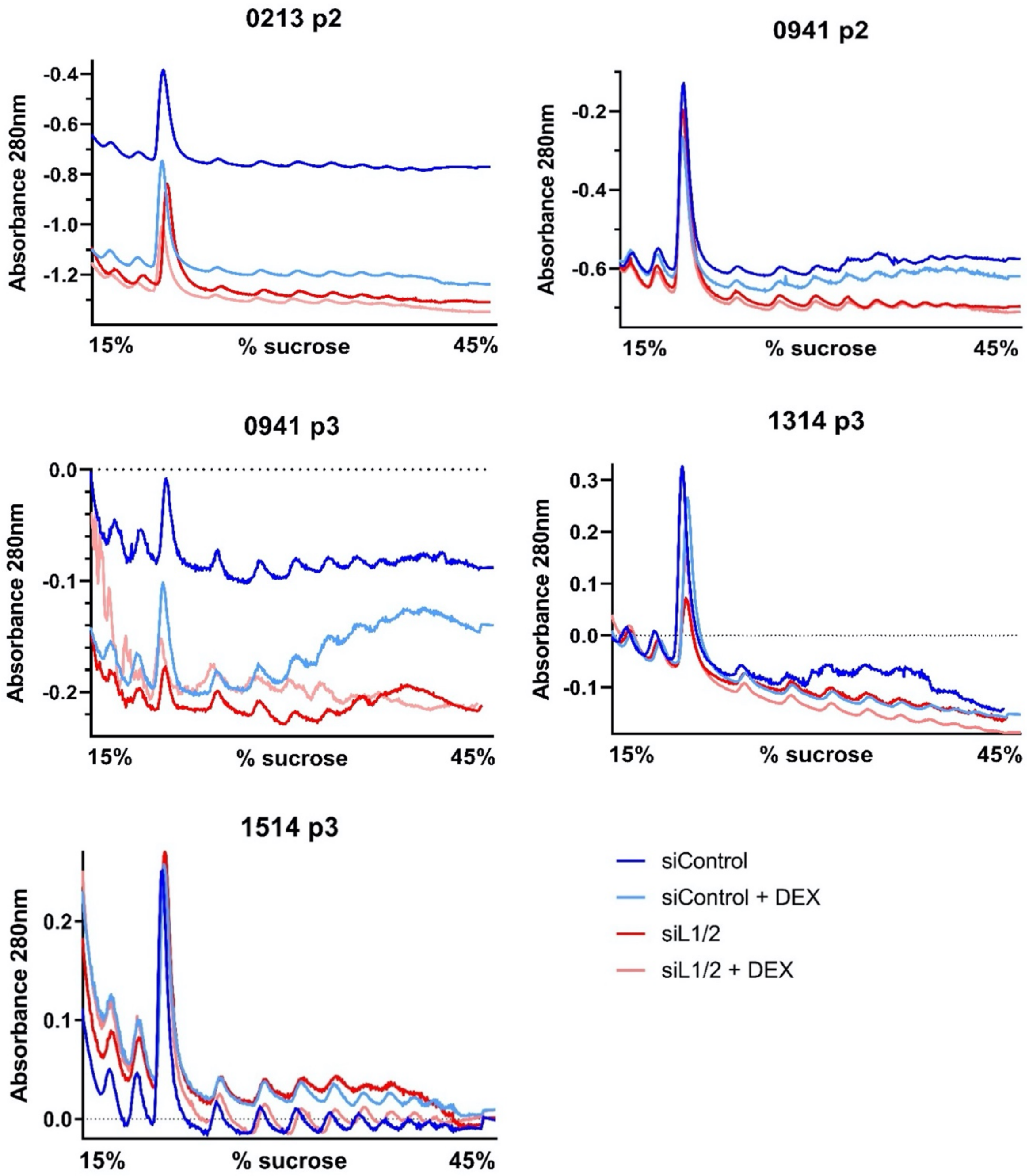


**Supplemental Figure 3. Polyribosome profiles from Frac-seq in primary BECs.**Polyribosome profiles from primary BECs employed in this study. Graphs depict the profiles from BECs from each donor (0213, 0941, 1314, 1514) and passage number (p2, p3). In successive order the peaks represent 40S, 60S, and 80S ribosomal subunits, and each subsequent peak two three, four, etc ribosomes together.

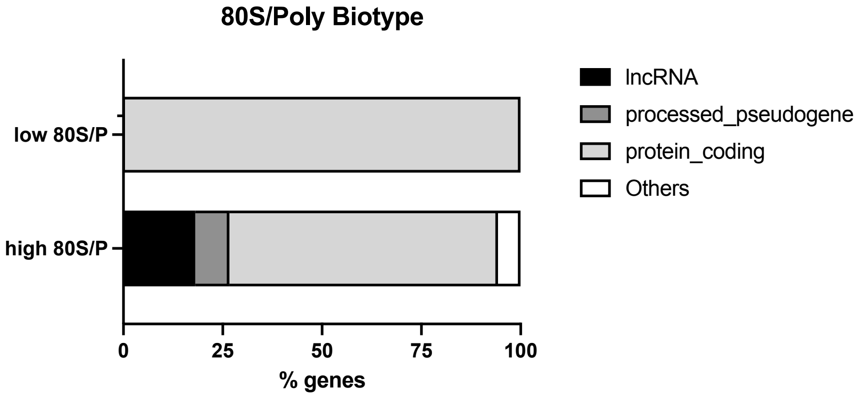


**Supplemental Figure 4. Differential RNA biotype associated to monosomes**. We analysed which transcripts were preferentially present in monosomes over their polyribosome detected levels. We firstly divided the baseMean counts from DESeq2 (in this case, the comparisons of dexamethasone effects in siControl transfected cells) to obtain a monosome/polyribosome ratio. We next filtered for genes that had more than 50 counts in monosomes, to increase the restriction in our analysis. We then sliced the bottom and top 10% candidates (low 80S/Poly, or *viceversa*) and analysed the biotype of RNA present in each population, plotting them in this bar graph. RNAs that had a low association to monosomes as compared to their polyribosome association were mainly protein coding genes (99.65%), while RNAs that had a high association to monosomes over polyribosomes had a large proportion of lncRNAs (18.07%) and processed pseudogenes (8.63%) with a drop of protein coding genes to 67.6%.

When the ratio of 80S/Total was analysed we found similar results (not shown) with protein coding genes in low 80S/Total being 99.65%, while RNAs that had a high association to monosomes over total had a large proportion of lncRNAs (17.48%) and processed pseudogenes (9.44%) with a drop of protein coding genes to 68.77%,

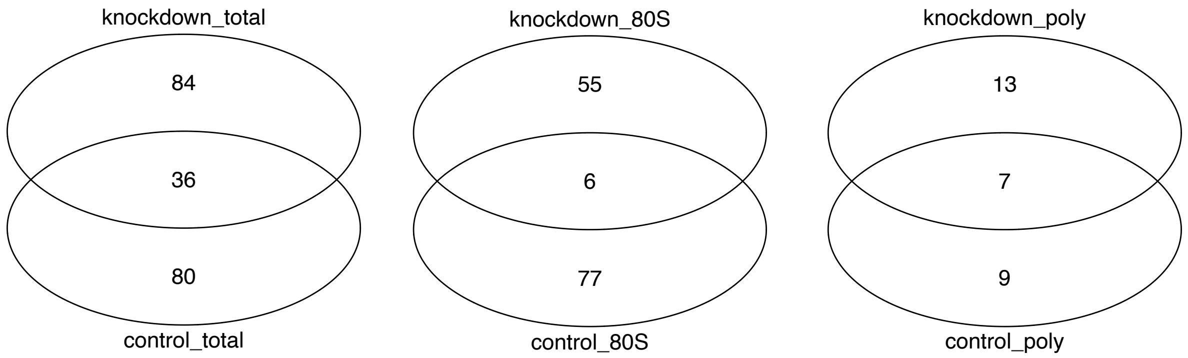


**Supplemental Figure 5. Overlap of GC modulated genes between siControl and siL1L2 depleted samples per fraction**. These Venn diagrams show the overlap of genes modulated by dexamethasone in siControl transfected samples (control) vs ZFP36L1/L2 depleted samples (knockdown) per fraction.
